# Modelling the influence of rhizodeposits on root water uptake

**DOI:** 10.1101/2025.03.28.645940

**Authors:** Andrew Mair, Emma Gómez Peral, Mariya Ptashnyk, Lionel Dupuy

## Abstract

The chemical compounds produced by plant roots, referred to generally as rhizodeposits, affect several soil hydraulic properties. For example, the surface tension of soil water, and the contact angle between menisci and the pore surface. What remains less clear is how these effects manifest when considering soil water infiltration and retention, and the consequent impact on the availability of water for uptake by plant roots. By modifying the Richards equation, a novel model for soil water transport was developed which incorporates the influences of rhizodeposits. The finite-element method was used for simulations of model equations and calibration against experimental data was achieved through Bayesian optimisation. Numerical simulations from the calibrated model were used to investigate the effects of rhizodeposits on root water uptake under various precipitation regimes. It was found that the impact of rhizodeposits on the availability of water to the roots can be either positive or negative depending on precipitation regime and root system maturity. This, therefore, suggests that rhizodeposit characteristics require careful consideration when developing crops for improved water use efficiency and stress-resilience.

## 1 Introduction

Improving plant water use efficiency is a central aim of agricultural research and, as previously outlined by Condon et al. (2004), there are, broadly speaking, 3 facets to this problem. The first is the ability of the plant to convert transpired water into biomass, the second is maximising the proportion of this biomass that then constitutes harvestable product, and the final aspect (the focus of this paper) is the root system’s capacity to access and uptake as much water as possible. An important characteristic to consider here is obviously the root system architecture (RSA), and, in this context, much scientific attention has been paid to ensuring that roots grow into soil regions with the highest water content. To this end, the most commonly promoted morphology is one where the majority of resources are allocated to growing downward so that stores of water deep in the soil can be accessed during periods of drought (Lynch, 2013; Uga et al., 2013; Lynch and Wojciechowski, 2015). Such root systems are less exposed to volatile rainfall patterns and, hence, more adapted to intensive agriculture (Wasson et al., 2012). RSA, however, not only determines where water can be extracted from but also influences the infiltration path of water following irrigation or rainfall (Ghestem et al., 2011; Noguchi et al., 1997, 1999). From this perspective, a desirable RSA is one that induces a pattern of infiltration in which the provision of water to the rooted zone is facilitated and losses of available water through evaporation and excessive drainage are minimised (Figure 1). Our previous study (Mair et al., 2023) showed how success in achieving this varied across a group of root systems with equal total biomass but contrasting architectural features.

**Figure 1:**
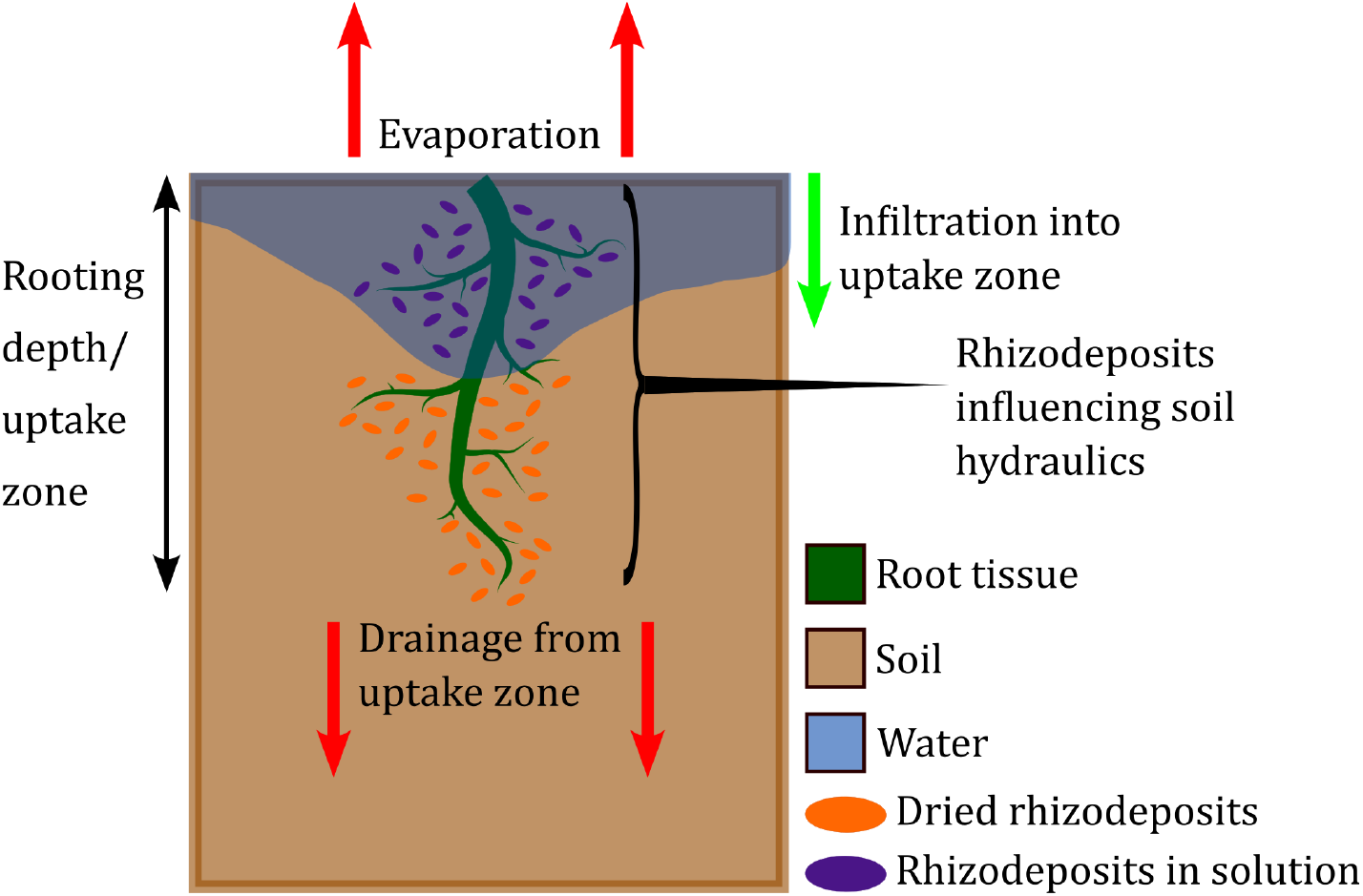
Rhizodeposits and root water uptake. The uptake performance of a plant is affected by the availability of water within the rooted regions of the soil. This figure shows the various processes that determine levels of water availability, all of which are influenced by the presence of rhizodeposits.

A root system’s architecture, though, is not the only trait that influences soil water dynamics. Depending on species, between 25% and 70% of the fixed carbon that is allocated by crops to their root systems is released into the soil in the form of rhizodeposits (McGrail et al., 2020). Across a range of crop types, these chemical compounds have been reported to affect a number of soil hydraulic properties, and the region falling under this influence is referred to as the rhizosphere. Specific observed effects include a decrease in the surface tension of the soil water solution as the concentration of wheat, maize or barley rhizodeposit is increased (Naveed et al., 2019; Read et al., 2003; Read and Gregory, 1997), and a reduction in its viscosity with higher levels of maize or barley rhizodeposit concentration (Naveed et al., 2019). Additionally, higher concentrations of sorbed maize and wheat rhizodeposits have been shown to increase the contact angle between the pore liquid and the soil surface (Ahmed et al., 2016; Benard et al., 2018; Naveed et al., 2019; Zickenrott et al., 2016). It is clear that all these rhizodeposit effects will impact differently on macroscopic water dynamics depending on the environmental conditions considered. However, knowledge of the consequences regarding the availability of water to the root system is lacking. This paper aims to show how the influence on water infiltration of the RSA and rhizodeposits of a given plant contributes to its water uptake efficiency. A mathematical model for water transport and root water uptake in vegetated soil is developed and then calibrated according to experimental observations for the effect of winter wheat exudates on water infiltration. Simulations from the calibrated model are then used to investigate the effect of increased rhizodeposit concentration on infiltration strength, evaporation levels and uptake performance. The effect is considered for root systems of different maturity and across environments that vary according to amounts of total precipitation and the pattern by which it is delivered.

## 2 Materials and Methods

### 2.1 A coupled model for soil water and rhizodeposit dynamics

Assuming a soil domain with depth *L* and spatial variable *x*, and a temporal interval with final time *T >* 0, the relationship between soil hydraulics, root rhizodeposits, and root water uptake is modelled by a system of equations that couples a modified Richards equation (Richards, 1931) solved for soil water pressure head *h* [L]:

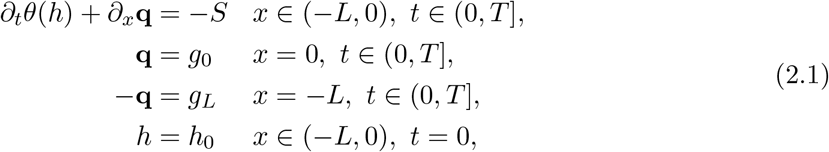

with an advection diffusion equation solved for the concentration of rhizodeposits *c*_*w*_ [ML^*−*3^] in the soil water solution:

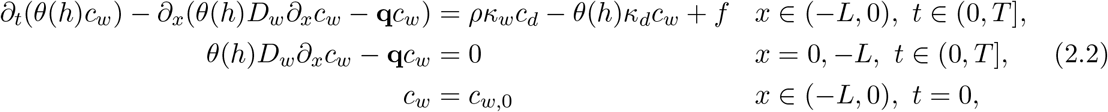

and an ordinary differential equation solved for the concentration of dried rhizodeposits attached to the soil particles *c*_*d*_ [MM^*−*1^]:

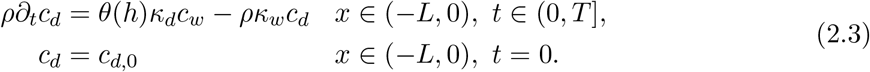

In (2.1), soil water content is given by *θ* [L^3^L^*−*3^], root water uptake by *S* [T^*−*1^], and the flux conditions at the upper surface and base of the soil by *g*_0_ and *g*_*L*_ respectively. The movement of water through the soil is driven by the Darcy-Buckingham flux (Darcy, 1856)

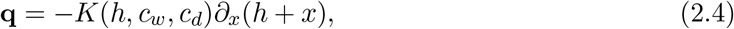

with *K* [LT^*−*1^] being the hydraulic conductivity. The flux **q** also determines the advective term in model (2.2) for the dynamics of solubilised rhizodeposits. The remaining terms in (2.2) and (2.3) are the rhizodeposit diffusion coefficient *D*_*w*_ [L^2^T^*−*1^], the soil bulk density *ρ* [ML^*−*3^], a source term *f* [ML^*−*3^T^*−*1^] for the release of rhizodeposits into the soil solution by the plant roots, and the rates at which solubilised rhizodeposits dry to the surface of the soil particles *κ*_*d*_ [T^*−*1^], and dried rhizodeposits solubilise *κ*_*w*_ [T^*−*1^].

The influence of rhizodeposits is incorporated into the water transport and uptake model (2.1) through modifications to the Van Genuchten (1980) and Mualem (1976) formulations of the functions for water content and hydraulic conductivity:

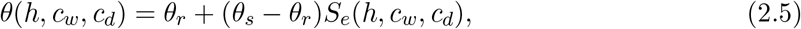

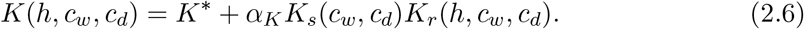

Here the effective soil saturation *S*_*e*_ in (2.5), the saturated hydraulic conductivity *K*_*s*_ [LT^*−*1^], and the effective hydraulic conductivity *K*_*r*_ in (2.6) are functions of pressure head and rhizodeposit concentration. The terms *θ*_*r*_ and *θ*_*s*_ in (2.5) give the residual and saturated soil water contents respectively and parameters *K*^*\**^ [LT^*−*1^] and *α*_*K*_ are included in (2.6) to account for the fact that water content and hydraulic conductivity values at a given pressure head are lower if a soil is wetting than if it is drying. This hysteresis exists because the processes that occur when a soil drains are generally governed by activity in smaller pores, whereas during wetting it is the activity within larger pores that is more consequential. Also, the contact angles between the menisci and the surface of soil particles are different depending on whether water is entering or draining from a pore (Van Genuchten and Pachepsky, 2011; Zhou, 2013). In (2.5) and (2.6) hysteresis is incorporated by the effective saturation as follows

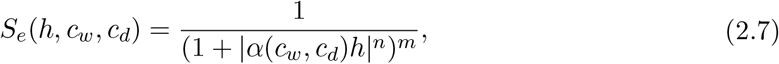

where *n* and 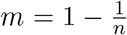 are shape parameters. For a depth *x* and time *t* the function *α* [*L*^*−*1^], relating to inverse pore air entry pressure, takes different values depending on whether the soil is wetting (increasing *h*) or drying (decreasing *h*):

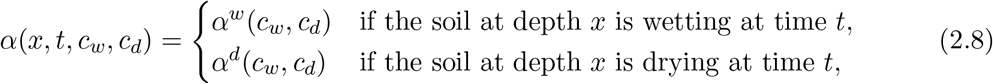

with *α*^*w*^ *≥ α*^*d*^ (Kool and Parker, 1987). Similarly, in (2.6) hysteresis is incorporated through the saturated hydraulic conductivity where:

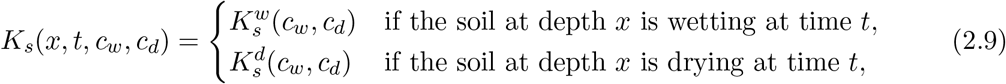

and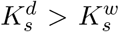 (Vogel et al., 1996). If at some depth and time (*x*_Δ_, *t*_Δ_) the soil changes from wetting to drying (or vice versa), then here *α* and *K*_*s*_ switch to *α*^*d*^ and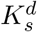 or *α*^*w*^ and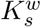 accordingly. To maintain the continuity of *θ* and *K* under transitions between wetting and drying, some other parameters must also change values. Specifically, with the pressure head, water content and hydraulic conductivity at the switch point (*x*_Δ_, *t*_Δ_) denoted as *h*_Δ_, *θ*_Δ_ and *K*_Δ_ respectively, the residual and saturated water contents assume the values

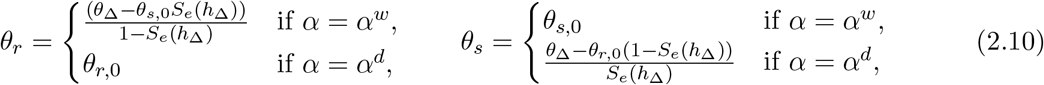

where *θ*_*r*,0_ and *θ*_*s*,0_ are the original non-hysteretic values of residual and saturated soil water content, and *K*^*\**^ and *α*_*K*_ take the values (Vogel et al., 1996):

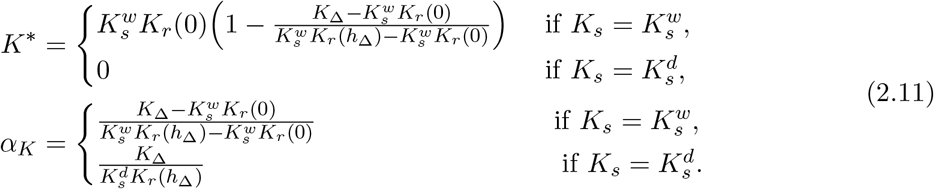

At *x*_Δ_, these new values (2.10)-(2.11) are then maintained until the next time that the soil at that depth switches from wetting to drying (or vice versa) and they have to be updated to new values using the same formulae. Finally, the expression for the effective hydraulic conductivity in (2.6) is:

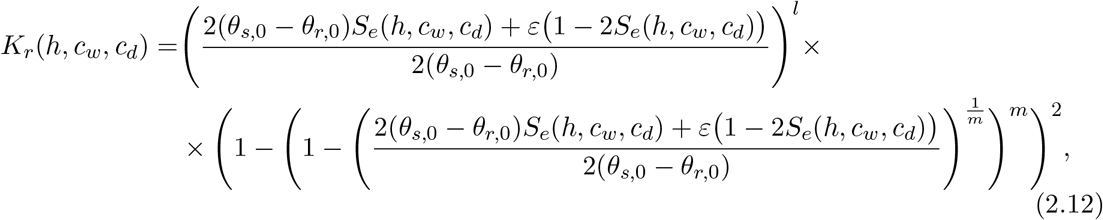

where *l* denotes the tortuosity of the porous structure (Mualem, 1976; Van Genuchten, 1980) and *ε >* 0 is a small regularisation parameter whose inclusion allows equations (2.1)-(2.3) to be solved numerically (List and Radu, 2016).

The surface tension of soil water *γ* and the contact angle *ϑ* of its menisci with pore surfaces have been shown to vary considerably according to the concentration of rhizodeposits, and the species of plant from which these were released (Ahmed et al., 2016; Naveed et al., 2018, 2019). The analysis here focuses on wheat plants where there is evidence that the rhizodeposits they release increase the soil-water contact angle (Zickenrott et al., 2016), and decrease the surface tension of the soil solution (Read et al., 2003). To account for this within *θ*, the approach of Karagunduz et al. (2001) is employed and scaling factors are introduced into *α*, such that for (*c*_*w*_, *c*_*d*_) *∈* [0, *∞*)^2^,

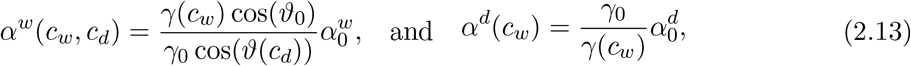

where *γ*_0_ and *ϑ*_0_ are the surface tension and contact angle values in the absence of rhizodeposits, and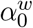 and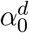 are the corresponding wetting and drying values of *α*. In the formulations of (2.13) a rhizodeposit-induced reduction in surface tension will decrease *α*^*w*^ and increase *α*^*d*^, which means the associated pressure head to a given water content will be decreased during soil wetting and increased during drying. Furthermore, an increase in contact angle will mean that the pressure head associated to a given water content during wetting is increased. In a similar way, to incorporate into *K* the effect of rhizodeposit-induced decrease in surface tension and increase in soil-water contact angle, the saturated hydraulic conductivities are formulated so that, for (*c*_*w*_, *c*_*d*_) *∈* [0, *∞*)^2^,

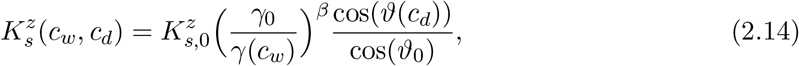

where *z* = *w, d*. This formulation of saturated hydraulic conductivity (2.14), was informed by the data of Gómez Peral et al. (2025) where it was found that solubilised winter wheat exudates facilitated the wetting of a transparent soil composed of Nafion™ particles. Moreover, this facilitation outweighed the soil hydrophobicity caused by some of the same exudates being dried onto the particles prior to the experiment. Assuming that a rhizodeposit-induced reduction in liquid surface tension was what caused the accelerated soil wetting, the exponent *β* in (2.14) determines the extent to which the infiltration facilitation from reduced-surface tension dominates over the hydrophobicity effect of increased contact angle.

As in (Simunek and Hopmans, 2009), root water uptake in (2.1) is modelled with a macroscopic sink term *S* incorporating water availability, root distribution, and potential plant transpiration *T*_p_ [*LT*^*−*1^]:

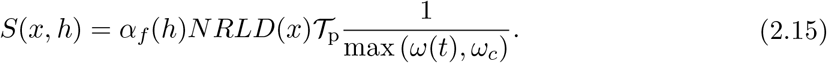

The dependence of uptake on the level of soil water content is included in *S* through the dimensionless stress response function 0 *≤α*_*f*_ *≤*1, and the plant’s capacity to reach the water within the soil is accounted for by the normalised root length density *NRLD* [*L*^*−*3^], which is a continuous function that gives the distribution of root length within the soil. The potential plant transpiration is formulated as *T*_p_ = ET_0_K_cb_, where ET_0_ [LT^*−*1^] is the reference evapotranspiration and K_cb_ [*−*] is the basal crop coefficient (Allen et al., 1998). The measure of the global water stress experienced by the plant is given by the function

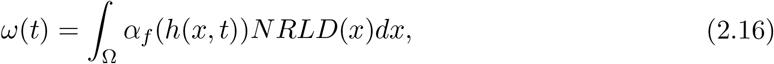

and the capacity of the plant to compensate for poor water availability in one soil region by increasing uptake in zones of higher saturation is reflected in the value of the critical water stress index *ω*_*c*_ (Cai et al., 2018). The boundary condition at the soil surface is

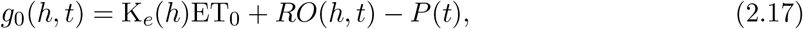

where K_*e*_ [*−*] is a function that determines the proportion of total evapotranspiration that comes from evaporation, *P* [LT^*−*1^] is the precipitation rate and *RO* [LT^*−*1^] is the run-off of water that occurs when precipitation cannot enter the soil due to the surface already being fully saturated:

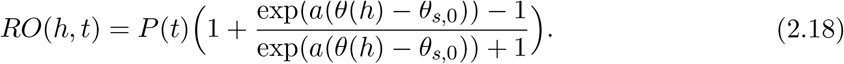

Here the constant *a >* 0 is set to a large enough value so that if the surface is fully saturated (*θ* = *θ*_*s*,0_) then *RO* = *P*. On the lower boundary *x* = *−L*, the condition of free drainage is assumed, which translates as *g*_*L*_(*h*) = *K*(*h, c*_*w*_, *c*_*d*_). Finally, the source term *f* in (2.2), for the release of rhizodeposits into the soil, is formulated as follows:

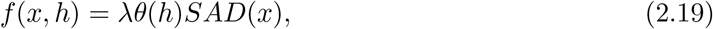

where *λ* [ML^*−*2^T^*−*1^] is the rate at which rhizodeposits are released from the surface of the root and *SAD* [L^*−*1^] is the root surface area density, a continuous function representing the distribution of root surface area throughout the soil.

### 2.2 Parametrisation and Calibration

The functions for normalised root length density *NRLD* and surface area density *SAD* were constructed for 3 winter wheat root systems of ages 6, 15 and 30 days. These root systems were generated from CRootBox (Schnepf et al., 2018) using the default settings for wheat systems and their respective architectures are shown in Figure 2. The methods previously employed and described in (Mair et al., 2022, 2023) were used to convert the architecture data of each root system into density functions for the distributions of root length and surface area. These functions took values in a 3D soil domain defined by the furthest reaching roots across each of the root systems, and were then integrated across the lateral dimensions to produce *NRLD* and *SAD* as functions of soil depth (*−L*, 0). Here *L* = 53 cm is the greatest depth reached by any root of the 3 systems. To explicitly model the ways in which rhizodeposits induce a reduction in soil water surface tension *γ*, and an increase in the contact angle *ϑ* between menisci and the soil pore surface, the data of Read et al. (2003) and Zickenrott et al. (2016) were used to obtain the following functions:

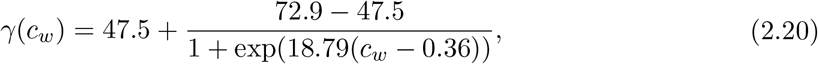

and

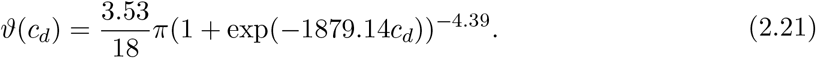

**Figure 2:**
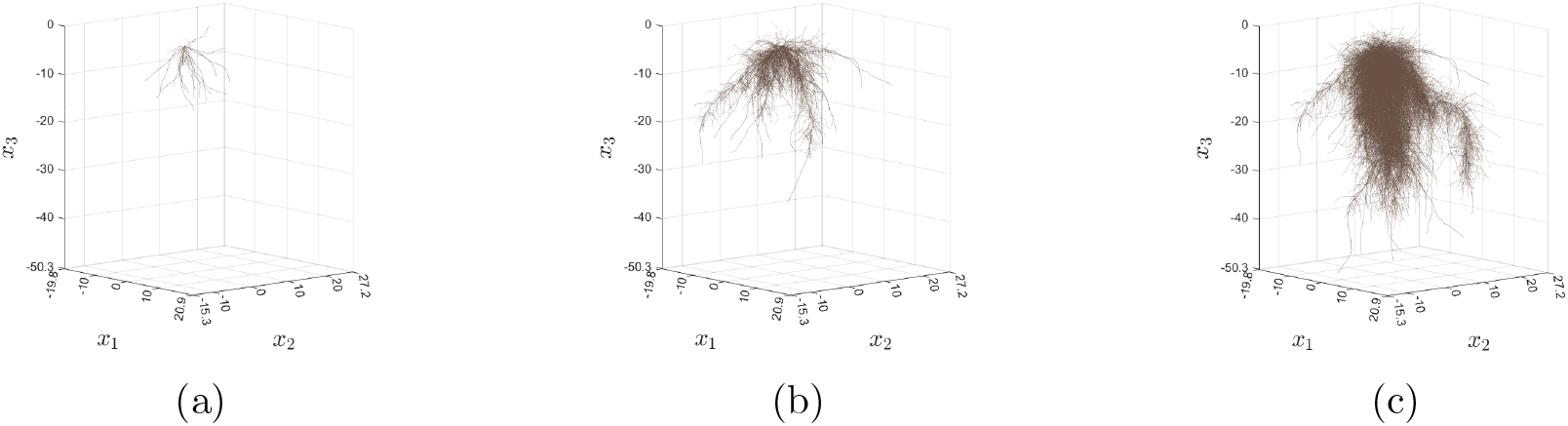
Simulated winter wheat root systems used for testing the impact of rhizodeposits on soil water dynamics and root water uptake. (a) 6-day old root system. (b) 15-day old root system. (c) 30-day old root system.

These functions return *γ*_0_ = *γ*(0) = 72.9 mNm^*−*1^ and *ϑ*_0_ = *ϑ*(0) = 0.0294 rad as the surface tension and contact angle when there are no rhizodeposits present. The precipitation term *P* is formulated as a step function of time *t* so that within chosen intervals of the simulation it prescribes constant rainfall at a given rate but outwith these it takes the value 0. A full description of the formulation of these type of functions is available in the supplementary material that accompanies (Mair et al., 2023).

A sandy loam soil was considered in all model simulations, and the majority of the values required to parametrise the models accordingly were taken from existing literature, see Table 1. The parameter values estimated from experimental data were the rate at which dried rhizodeposits solubilise *κ*_*w*_, the rate at which solubilised rhizodeposits dry to the pore surface *κ*_*d*_, and *β* the extent to which a rhizodeposit-induced reduction in surface tension increases hydraulic conductivity. These 3 values were approximated by Gómez Peral et al. (2025) who used a dye tracing experiment (Liu et al., 2025), in which chambers were filled with 3 layers of the artificial transparent soil Nafion™, and the infiltration of dye solution through the medium was monitored. Extracted winter wheat rhizodeposits were added to the dye solution in the upper Nafion layer™ and the effect on infiltration was recorded. The model (2.1)-(2.3), and an advection-diffusion equation for dye concentration within the chamber, were then formulated to simulate the process, and Bayesian optimisation (Brochu et al., 2010) was used to approximate the values (*κ*_*d*_, *κ*_*w*_, *β*) that gave this model the capacity to most accurately reproduce the experimental results. These values are also shown in Table 1.

**Table 1:** Notation and description of terms that appear in this article. The value column reads n/a if the corresponding term is a variable, a function, or a parameter that is altered between simulations. References included indicate if the parameter value or function formulation were taken from a specific source.

| Term | Description | Value | Reference |
| --- | --- | --- | --- |
| $a$ | Steepness of transition to complete run off once the boundary approaches saturation | 250 | n/a |
| $c_d$ | Concentration of rhizodeposits dried to soil surface ($\text{mgmg}^{-1}$) | n/a | n/a |
| $c_{d,0}$ | Initial concentration of rhizodeposits dried to soil surface ($\text{mgmg}^{-1}$) | n/a | n/a |
| $c_w$ | Concentration of rhizodeposits in solution ($\text{mgcm}^{-3}$) | n/a | n/a |
| $c_{w,0}$ | Initial concentration of rhizodeposits in solution ($\text{mgcm}^{-3}$) | n/a | n/a |
| $D_w$ | Rhizodeposit diffusion coefficient ($\text{cm}^2\text{d}^{-1}$) | 0.65 | (Scott et al., 1995) |
| $\text{ET}_0$ | Reference evapotranspiration ($\text{cmd}^{-1}$) | 0.1 | (Allen et al., 1998) |
| $f$ | Rhizodeposit release rate ($\text{mgcm}^{-3}\text{d}^{-1}$) | n/a | n/a |
| $f_c$ | Fraction of soil surface covered by vegetation (-) (appears in $K_e$) | 0.1 | (Allen et al., 1998, Chapter 7, Table 21) |
| $f_w$ | Fraction of soil surface wetted by precipitation (-) (appears in $K_e$) | 1.0 | (Allen et al., 1998, Chapter 7, Table 20) |
| $g_0$ | Water flux at upper soil surface ($\text{cmd}^{-1}$) | n/a | n/a |
| $gL$ | Water flux at lower soil surface ($\text{cmd}^{-1}$) | n/a | n/a |
| $h$ | Soil water pressure head (cm) | n/a | n/a |
| $h_0$ | Initial pressure head condition (cm) | n/a | n/a |
| $h_1$ | Pressure head associated with anaerobiosis due to soil saturation (cm) (appears in $\alpha_f$) | 0 | (Wesseling, 1991, Table 5) |
| $h_2$ | Upper limit of pressure head interval in which uptake capacity is maximised (cm) (appears in $\alpha_f$) | -1.0 | (Wesseling, 1991, Table 5) |
| $h_3$ | Lower limit of pressure head interval in which uptake capacity is maximised (cm) (appears in $\alpha_f$) | -500.0 | (Wesseling, 1991, Table 5) |
| $h_4$ | Pressure head associated with plant wilting point (cm) (appears in $\alpha_f$) | -16000.0 | (Wesseling, 1991, Table 5) |
| $h_\Delta$ | Pressure head at hysteresis reversal point (cm) | n/a | (Kool and Parker, 1987) |
| $K$ | Hydraulic conductivity ($\text{cmd}^{-1}$) | n/a | (Mualem, 1976; Van Genuchten, 1980) |
| $K_e$ | Evaporation coefficient (-) | n/a | (Allen et al., 1998; Mair et al., 2023) |
| $K_r$ | Effective hydraulic conductivity (-) | n/a | (Mualem, 1976; Van Genuchten, 1980) |
| $K_s$ | Hysteretic saturated hydraulic conductivity ($\text{cmd}^{-1}$) | n/a | (Vogel et al., 1996) |
| $K_s^d$ | Saturated hydraulic conductivity of drying hydraulic conductivity curve ($\text{cmd}^{-1}$) | n/a | (Vogel et al., 1996) |
| $K_{s,0}^d$ | Saturated hydraulic conductivity of drying hydraulic conductivity curve in absence of rhizodeposits ($\text{cmd}^{-1}$) | 120 | (Vogel et al., 1996) |
| $K_s^w$ | Saturated hydraulic conductivity of wetting hydraulic conductivity curve ($\text{cmd}^{-1}$) | n/a | (Vogel et al., 1996) |
| $K_{s,0}^w$ | Saturated hydraulic conductivity of wetting hydraulic conductivity curve in absence of rhizodeposits ($\text{cmd}^{-1}$) | 72 | (Vogel et al., 1996) |
| $K_\Delta$ | Hydraulic conductivity at hysteresis reversal point ($\text{cmd}^{-1}$) | n/a | (Vogel et al., 1996) |
| $K^*$ | Fictitious hysteresis parameter in hydraulic conductivity ($\text{cmd}^{-1}$) | n/a | (Vogel et al., 1996) |
| $K_{cb}$ | Basal crop coefficient (-) | n/a | (Allen et al., 1998) |
| $K_{c\max}$ | Maximum crop coefficient (-) (appears in $K_e$) | 1.30 | (Allen et al., 1998, Chapter 7, Table 19) |
| $l$ | Pore tortuosity parameter in hydraulic conductivity function (-) | 0.5 | (Van Genuchten and Pachepsky, 2011) |
| $L$ | Soil depth (cm) | 53 | n/a |
| $[L]$ | Length dimension | n/a | n/a |
| $m$ | Pore size parameter of soil water retention curve | 0.47 | (Mualem, 1976; Van Genuchten, 1980) |
| $[M]$ | Mass dimension | n/a | n/a |
| $n$ | Pore size parameter of soil water retention curve | 1.89 | (Mualem, 1976; Van Genuchten, 1980) |
| $NRLD$ | Normalised root length density ($\text{cm}^{-3}$) | n/a | (Mair et al., 2023) |
| $P$ | Precipitation rate ($\text{cmd}^{-1}$) | n/a | n/a |
| $\mathbf{q}$ | Soil water flux ($\text{cmd}^{-1}$) | n/a | n/a |
| $RO$ | Surface runoff ($\text{cmd}^{-1}$) | n/a | n/a |
| $S$ | Root water uptake rate ($\text{cmd}^{-1}$) | n/a | (Simunek and Hopmans, 2009) |
| $S_e$ | Effective saturation (-) | n/a | (Mualem, 1976; Van Genuchten, 1980) |
| $SAD$ | Root surface area density ($\text{cm}^{-1}$) | n/a | (Mair et al., 2023) |
| $t$ | Temporal variable (d) | n/a | n/a |
| $T$ | Final time (d) | 3 | n/a |
| $[T]$ | Time dimension | n/a | n/a |
| $\mathcal{T}_p$ | Potential plant transpiration ($\text{cmd}^{-1}$) | n/a | (Allen et al., 1998) |
| $x$ | Spatial variable (cm) | n/a | n/a |
| $\alpha$ | Inverse air entry pressure head ($\text{cm}^{-1}$) | n/a | (Kool and Parker, 1987) |
| $\alpha_f$ | Water stress response (-) | n/a | (Simunek and Hopmans, 2009) |
| $\alpha_K$ | Hydraulic conductivity hysteresis scaling parameter ($\text{cmd}^{-1}$) | n/a | (Vogel et al., 1996) |
| $\alpha^d$ | Value of $\alpha$ in drying branch of soil water retention curve ($\text{cm}^{-1}$) | n/a | (Kool and Parker, 1987) |
| $\alpha_0^d$ | Non-hysteretic value of $\alpha$ in drying soil water retention curve in absence of rhizodeposits ($\text{cmd}^{-1}$). | 0.0114 | (Carsel and Parrish, 1988) |
| $\alpha^w$ | Value of $\alpha$ in wetting branch of soil water retention curve ($\text{cm}^{-1}$) | n/a | (Kool and Parker, 1987) |
| $\alpha_0^w$ | Value of $\alpha$ in wetting soil water retention curve in absence of rhizodeposits ($\text{cmd}^{-1}$). | 0.0521 | (Carsel and Parrish, 1988) |
| $\beta$ | Fitting parameter in $K_s^w$ and $K_s^d$ (-) | 8.772 | (Gómez Peral et al., 2025) |
| $\varepsilon$ | Regularisation parameter in effective hydraulic conductivity (-) | $10^{-15}$ | n/a |
| $\gamma$ | Surface tension of soil water ($\text{mNm}^{-1}$) | n/a | n/a |
| $\gamma_0$ | Value of $\gamma$ in absence of rhizodeposits ($\text{mNm}^{-1}$). | 72.9 | (Read et al., 2003) |
| $\kappa_d$ | Rhizodeposit drying rate ($\text{d}^{-1}$) | $9.94 \times 10^{-2}$ | (Gómez Peral et al., 2025) |
| $\kappa_w$ | Rhizodeposit wetting rate ($\text{d}^{-1}$) | $1.78 \times 10^{-2}$ | (Gómez Peral et al., 2025) |
| $\lambda$ | Rate of rhizodeposit release per unit root surface area ($\text{mgcm}^{-2}\text{d}^{-1}$) | 0.039 | (Ptashnyk et al., 2011) |
| $\rho$ | Soil bulk density ($\text{mgcm}^{-3}$) | 1650 | (Morris and Lowery, 1988) |
| $\theta$ | Soil water content ($\text{cm}^3\text{cm}^{-3}$) | n/a | (Mualem, 1976; Van Genuchten, 1980) |
| $\theta_{fc}$ | Water content at field capacity ($\text{cm}^3\text{cm}^{-3}$) (appears in $K_e$) | 0.28 | (Allen et al., 1998, Chapter 7, Table 19) |
| $\theta_r$ | Hysteretic residual water content ($\text{cm}^3\text{cm}^{-3}$) | n/a | (Kool and Parker, 1987) |
| $\theta_{r,0}$ | Non-hysteretic residual water content ($\text{cm}^3\text{cm}^{-3}$) | 0.065 | (Carsel and Parrish, 1988) |
| $\theta_s$ | Hysteretic saturated water content ($\text{cm}^3\text{cm}^{-3}$) | n/a | (Kool and Parker, 1987) |
| $\theta_{s,0}$ | Non-hysteretic saturated water content ($\text{cm}^3\text{cm}^{-3}$) | 0.41 | (Carsel and Parrish, 1988) |
| $\theta_{wp}$ | Water content at wilting point ($\text{cm}^3\text{cm}^{-3}$) (appears in $K_e$) | 0.16 | (Allen et al., 1998, Chapter 7, Table 19) |
| $\theta_\Delta$ | Water content at hysteresis reversal point ($\text{cm}^3\text{cm}^{-3}$) | n/a | (Kool and Parker, 1987) |
| $\vartheta$ | Contact angle between soil water and pore surface (rad) | n/a | n/a |
| $\vartheta_0$ | Value of $\vartheta$ in absence of rhizodeposits (rad). | 0.0294 | (Zickenrott et al., 2016) |
| $\omega$ | Global water stress ($\text{cm}^{-2}$) | n/a | (Simunek and Hopmans, 2009) |
| $\omega_c$ | Critical water stress index ($\text{cm}^{-2}$) | 0.8 | (Cai et al., 2018) |

### 2.3 Simulated scenarios

To investigate the impact of rhizodeposits on root water uptake performance, the model system (2.1)-(2.3) was parametrised with different *NRLD* and *SAD* functions for each root system age as well as varying basal crop coefficients K_cb_. Specifically, data from (Allen et al., 1998) was used to formulate a piecewise linear function of K_cb_ with respect to age, and this resulted in values of K_cb_ = 0.15, 0.32, 0.5 for the 6, 15 and 30-day old root systems respectively. For every root system parametrisation, 6 precipitation regimes were considered. In 3 of these regimes the precipitation function *P* was formulated to deliver a total of 0.12 cm split over either 1, 2 or 3 rainfall events within the time period *T* = 3 d. The remaining 3 precipitation regimes were defined in the same way, but with total rainfall set to 0.28 cm. In each scenario, initial water content was set to a uniform value of *θ*(*h*_0_) = 0.69 throughout the domain. In the scenarios that included rhizodeposits, the initial concentration of rhizodeposit in solution was set so that in the region of the soil occupied by roots *c*_*w*,0_ = 2.2 mgcm^*−*3^ and in the part of the domain without roots *c*_*w*,0_ = 0 mgcm^*−*3^. Assuming an initial equilibrium between the dynamics of dried rhizodeposits and rhizodeposits in solution, initial dried rhizodeposit concentration was set so that *c*_*d*,0_ = 5.1 *×* 10^*−*4^ mgcm^*−*3^ in the part of the soil with roots present, and *c*_*d*,0_ = 0 mgcm^*−*3^ in the part without. Water infiltration simulations were then obtained for each combination of root system and precipitation pattern, both with and without the incorporation of rhizodeposit influence, and results on soil water dynamics and root water uptake were recorded.

### 2.4 Computations

Numerical solutions for pressure head, rhizodeposits in solution and dried rhizodeposits were obtained from the model (2.1)-(2.3) using a conformal finite element method and an implicit Euler discretisation in time. The intervals of the 1D domain discretisation were each 0.593 cm in length, and the time step size was 0.001 d. Linearisation of the functions for water content and hydraulic conductivity in (2.1) was carried out using the L-scheme of List and Radu (2016). The numerical method was implemented using the FEniCS library (Alnæs et al., 2015) and the algorithms to construct *NRLD* and *SAD* from CRootBox architectures were taken from (Mair et al., 2022, 2023) and run in Python 3 using the libraries NumPy and SciPy (Harris et al., 2020). The visualisations of 1D root density profiles, water distributions and flux strength were generated in Paraview (Ahrens et al., 2005). The plots in Figure 2, showing the architectures of the studied root systems, were generated using code of Dupuy et al. (2005), which was run in MATLAB 2024b. The Bayesian optimisation algorithm used to calibrate model (2.1)-(2.3) was implemented using the scikit-optimise library in Python 3 (Head et al., 2018).

## 3 Results

### 3.1 Effect of rhizodeposits on water infiltration

In the case where total precipitation of 0.12 cm was shared over 3 rainfall events, simulations showed that water infiltration was greater in all scenarios where rhizodeposits were present compared to when they were not (Figure 3). Water infiltration increased with rainfall both with and without the presence of rhizodeposits. However, the increased infiltration in the presence of rhizodeposits became more and more pronounced with each rainfall event (Figure 3). At all observed time points, and for each age of root system, this increased water infiltration in the presence of rhizodeposits lead to a higher water content at the soil depth where root length density was highest (Figure 3). Moreover, at no point of the simulation did this enhanced water infiltration appear to have caused much movement of water into deeper non-rooted soil layers. In the scenario where a total precipitation of 0.28 cm was delivered in one day (Figure 4), water infiltration was stronger than in the case where the delivery of a total precipitation of 0.12 cm was split over 3 events (Figure 3). Like with 0.12 cm precipitation delivered in 3 events, however, when 0.28 cm was delivered in one day, the water infiltration was always stronger with rhizodeposits present than without (Figure 4). Similarly true for both cases was that at 0.5 days, just after the single/first rainfall event, the increased infiltration of water had resulted in an increased water content in the depths of soil where the root length density of all root systems was highest. In the case of 0.12 cm delivered in 3 events, the strength of this water infiltration increased over the simulation (Figure 3). In contrast, though, with 0.28 cm delivered in one day, the infiltration strength was greatest at 0.5 days and then decreased over the rest of the simulation (Figure 4). Furthermore, for the 6 and 15-day old root systems with rhizodeposits present and 0.28 cm precipitation delivered in one event, the increase in infiltration was excessive and after 1.5 days had already resulted in a considerable amount of water being lost to depths with low or zero root length density (Figure 4). Since the roots of the 30-day old root system were allocated more equally and to greater depths of the soil, this increased infiltration did not reduce the availability of water for uptake in the same way that it did for the younger plants.

**Figure 3:**
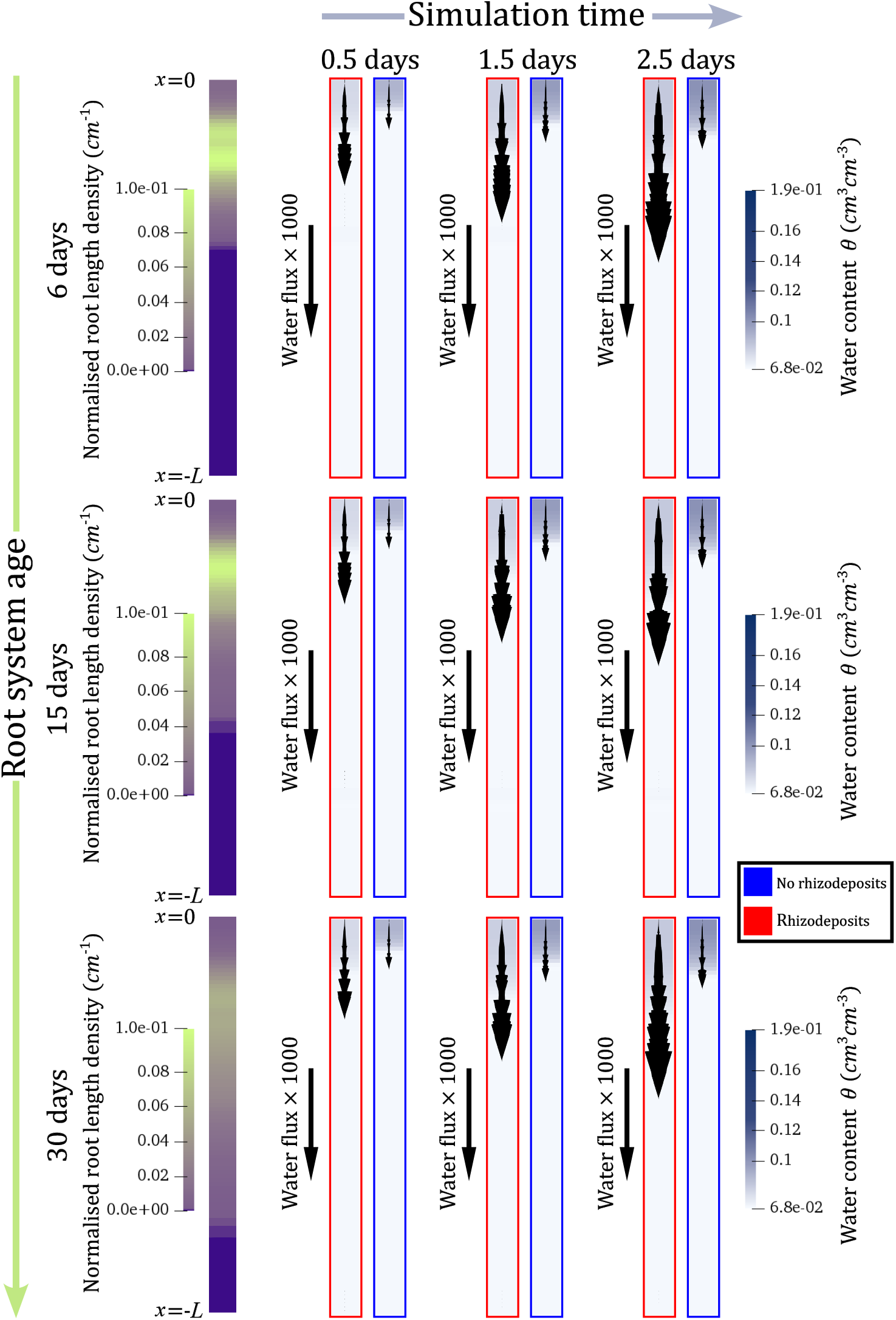
The influence of rhizodeposits on the evolution of soil water content and flux over time for the regime in which 0.12 cm total rainfall is split over 3 events. The figures of the leftmost column show the normalised root length density profiles of each plant within the 1D soil domain (age increasing downwards). The remaining columns show snapshots of the water dynamics within the domain at different time points of the simulation. Each row relates to soil vegetated by the roots of each age of plant, and the arrows illustrate the strength of the flow at each time point.

**Figure 4:**
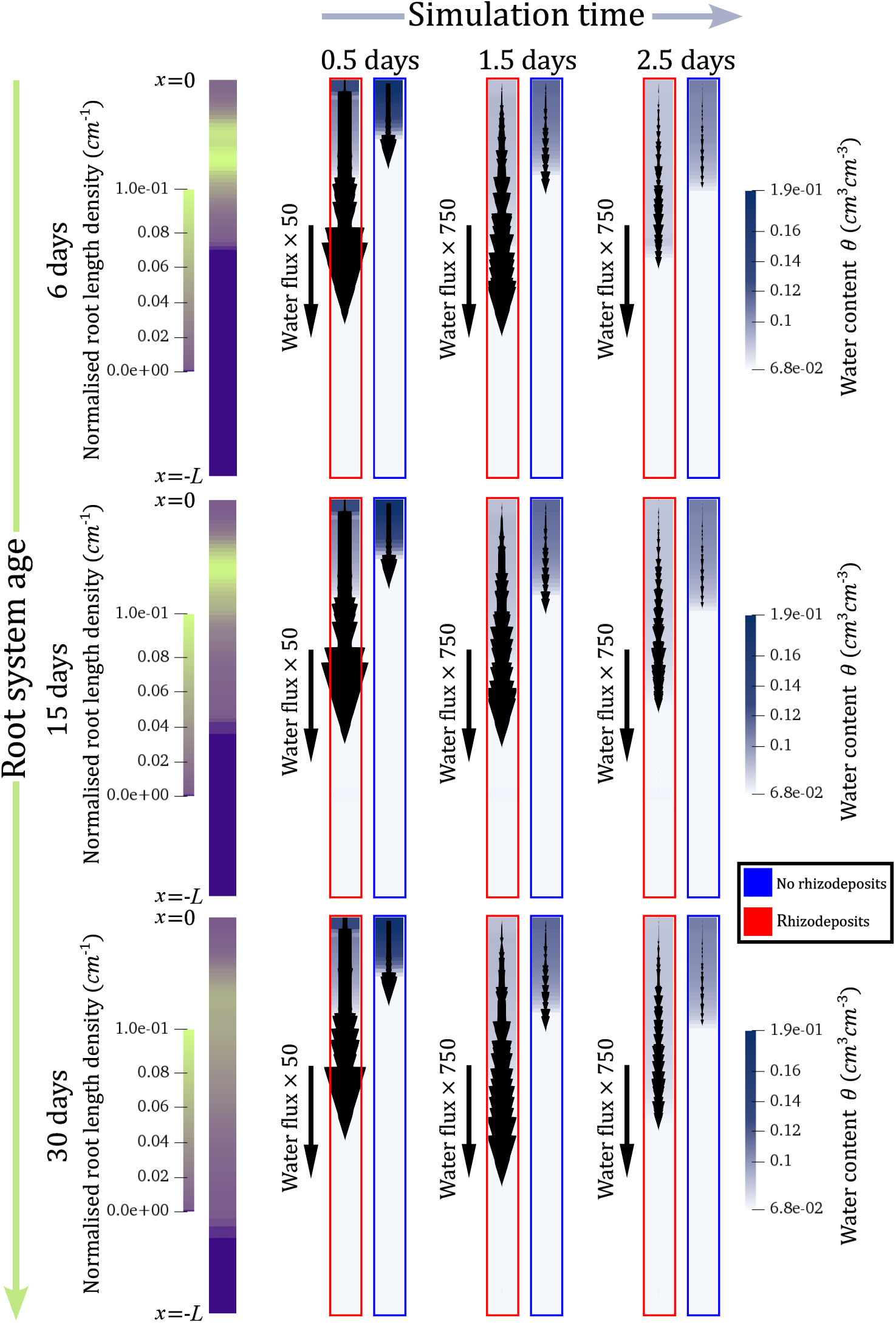
The influence of rhizodeposits on the evolution of soil water content and flux over time for the regime in which 0.28 cm of rainfall is delivered in one event. The figures of the leftmost column show the normalised root length density profiles of each plant within the 1D soil domain (age increasing downwards). The remaining columns show snapshots of the water dynamics within the domain at different time points of the simulation. Each row relates to soil vegetated by the roots of each age of plant, and the arrows illustrate the strength of the flow at each time point.

### 3.2 Effect of rhizodeposits on rates of soil surface evaporation and root water uptake

For all root systems in all precipitation conditions, evaporation rates were lower, at every point of the 3 day simulation, with rhizodeposits present in the soil than without (Figures 5 and 6 (d)-(f)). It was also the case for all precipitation regimes that, following a rainfall event, the water uptake rates of all root systems in soils with rhizodeposits became larger (for at least a finite time period) than those in soils without rhizodeposits (Figures 5 and 6 (g)-(i)). For the 30-day old root system, which had the deepest rooting depth, this higher uptake rate in the presence of rhizodeposits was observed at all points in time, and this was true across all precipitation regimes (Figures 5 and 6 (g)-(i)). Moreover, at all time points of the regimes with lower total precipitation (0.12 cm), the uptake rates of the 6 and 15-day old plants, which had shallower root architectures, were also higher with rhizodeposits present than without (Figure 5). However, it can be seen that in the precipitation pattern where all 0.12 cm was delivered in a single event, the difference between the increased uptake rate of the 6-day plant with rhizodeposits and the lower rate without rhizodeposits had almost disappeared by the end of the simulation (Figure 5 (g)). This, therefore, suggested that a higher uptake rate with rhizodeposits than without may not necessarily be maintained for the entire simulation under all combinations of total precipitation, delivery pattern and root system age. Indeed, under all regimes with 0.28 cm of total precipitation the shallower 6 and 15-day old root systems initially assumed higher uptake rates with rhizodeposits in the soil but, eventually, the uptake rates in soil without rhizodeposits overtook and maintained higher values until the end of the simulation (Figure 6). The timing of this overtake was earliest for the precipitation pattern in which all 0.28 cm of water is delivered in one event.

**Figure 5:**
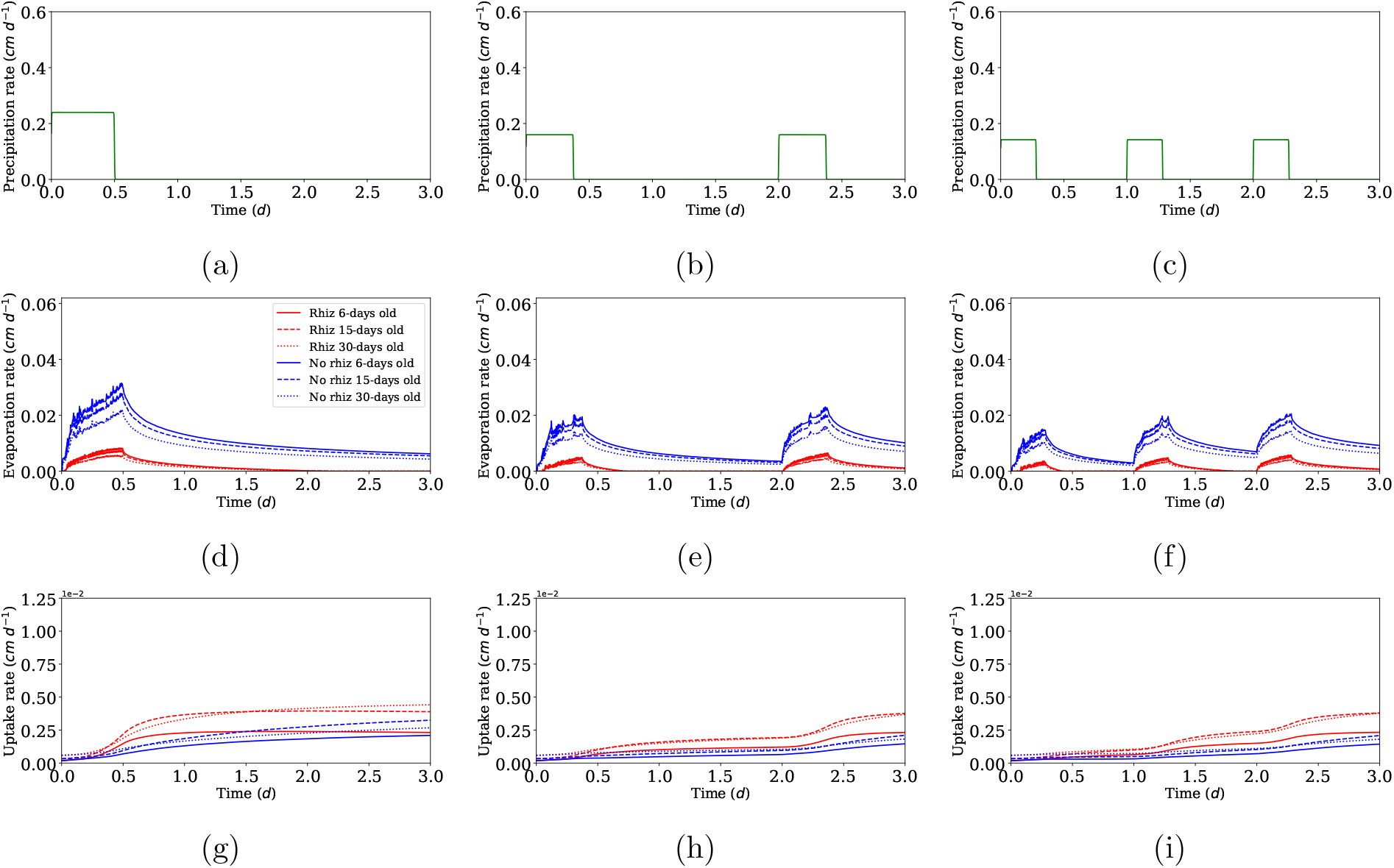
The influence of rhizodeposits on root water uptake under the different delivery patterns of lower total precipitation (0.12 cm). Plots (a)-(c) show the precipitation patterns considered. The plots (d)-(f) show the evaporation rates corresponding to each pattern and for soils occupied by each age of root system with or without rhizodeposits present. Similarly plots (g)-(i) show the corresponding uptake rates of each root system, with and without rhizodeposits present.

**Figure 6:**
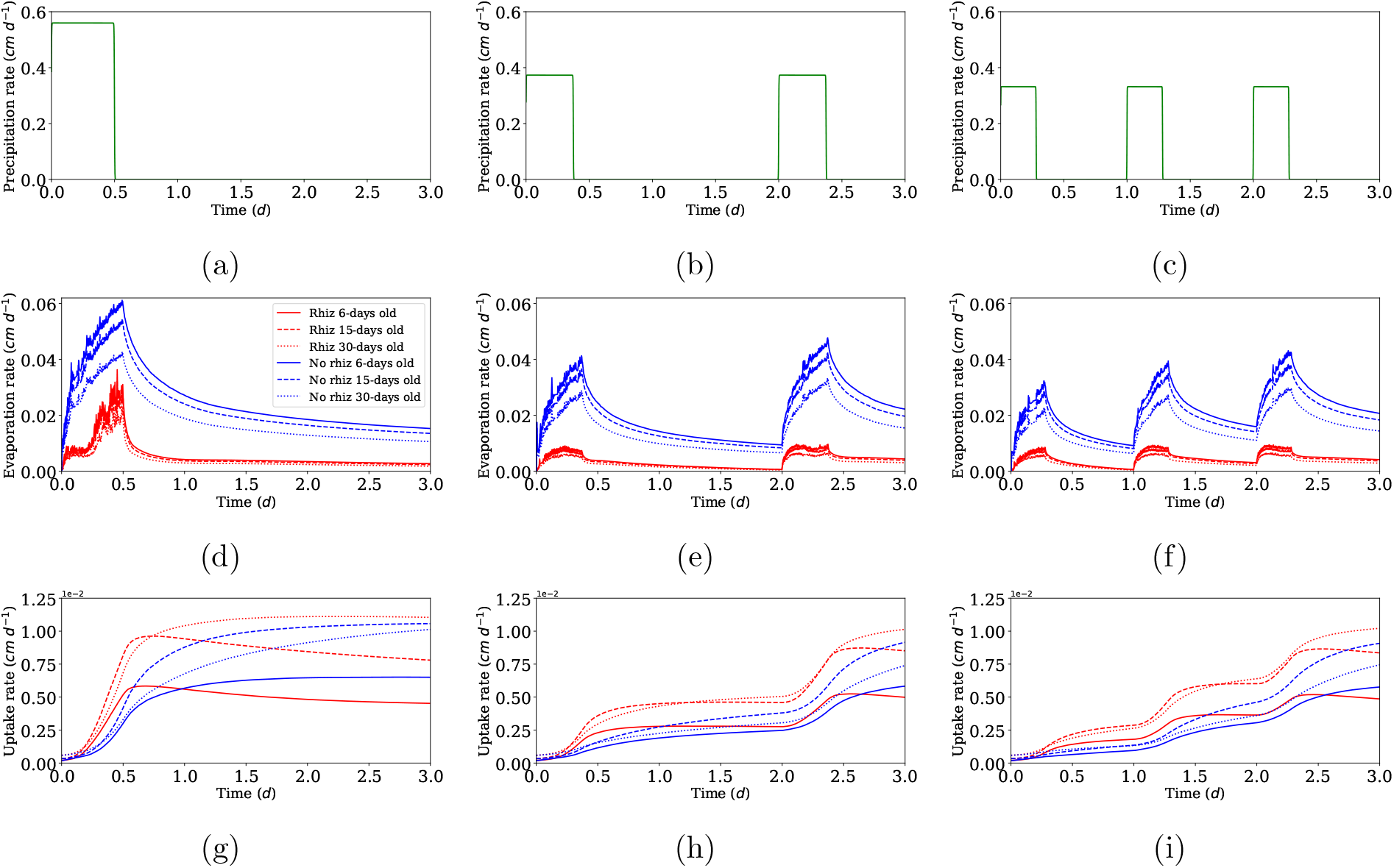
The influence of rhizodeposits on root water uptake under the different delivery patterns of higher total precipitation (0.28 cm). Plots (a)-(c) show the precipitation patterns considered. The plots (d)-(f) show the evaporation rates corresponding to each pattern and for soils occupied by each age of root system with or without rhizodeposits present. Similarly plots (g)-(i) show the corresponding uptake rates of each root system, with and without rhizodeposits present.

Finally, the last relevant finding regarding uptake rates was that within certain time intervals of these 3 day simulations (if not the entire simulation) the uptake rates of the 15-day old root system were slightly higher than those of the 30-day old, and this was true for soils with or without rhizodeposits (Figures 5 and 6). This resulted from an effect of using a macroscopic uptake function based on normalised root length density to compare performance of root systems of different ages. To our knowledge, this effect has not been previously reported and, as such, will be explained further in the discussion.

### 3.3 Rhizodeposit effect on total uptake and the sensitivity to precipitation pattern and root system maturity

In all rainfall patterns with 0.12 cm total precipitation, total water uptake of every age of root system was greater if rhizodeposits were present in the soil than if they were not (Table 2). Moreover, for all rainfall patterns of 0.28 cm total precipitation the total uptake of the 30-day old root system was also always larger when rhizodeposits were present than when they were not (Table 3). These results are expected given that, as previously mentioned, in these cases the higher uptake rates in the presence of rhizodeposits were maintained over the entire simulation period. The more nuanced results come from the cases where uptake rates with rhizodeposits in the soil were at some stage overtaken by the uptake rates without rhizodeposits. Specifically, in the regimes with a total precipitation of 0.28 cm, if it was delivered in 2 or 3 rainfall events, then the 6 and 15-day old root systems with rhizodeposits also continued to have higher total root water uptake than without rhizodeposits (Table 2). However, when the entire 0.28 cm was delivered in one event, the uptake rates of the 6 and 15-day old root systems in soil with rhizodeposits were overtaken earlier in the simulation by the uptake rates without rhizodeposits, and their total uptake ended up being lower with rhizodeposits in the soil than without (Table 3). This result can also be seen in Figure 7 which shows that the percentage change in total water uptake resulting from the presence of rhizodeposits was negative in the case where 0.28 cm of precipitation was delivered in one event and the root systems were 6 or 15 days old. Considering root systems of all ages, there was in fact a general trend that the percentage positive change in total water uptake from including rhizodeposits in the simulations decreased as the amount of total rainfall in the 3 day period was increased (Figure 7). In addition, when fixing total rainfall and considering the pattern by which it was delivered, it is also true that the positive effect of rhizodeposits on total water uptake was reduced as the precipitation was delivered over fewer events (Figure 7).

**Table 2:** Total uptake of each root system with and without rhizodeposits for each rainfall distribution of the lower-rainfall regime.

| Total rainfall | 0.12 cm |  |  |  |  |  |
| --- | --- | --- | --- | --- | --- | --- |
| Rainfall distribution | 1 event |  | 2 events |  | 3 events |  |
| Rhizodeposits | No | Yes | No | Yes | No | Yes |
| Total uptake: 6-day old plant (cm) | $4.31 \times 10^{-3}$ | $6.04 \times 10^{-3}$ | $1.99 \times 10^{-3}$ | $3.64 \times 10^{-3}$ | $1.91 \times 10^{-3}$ | $3.64 \times 10^{-3}$ |
| Total uptake: 15-day old plant (cm) | $6.39 \times 10^{-3}$ | $9.83 \times 10^{-3}$ | $2.89 \times 10^{-3}$ | $5.75 \times 10^{-3}$ | $2.78 \times 10^{-3}$ | $5.77 \times 10^{-3}$ |
| Total uptake: 30-day old plant (cm) | $5.58 \times 10^{-3}$ | $10.02 \times 10^{-3}$ | $3.16 \times 10^{-3}$ | $5.59 \times 10^{-3}$ | $3.08 \times 10^{-3}$ | $5.63 \times 10^{-3}$ |

**Table 3:** Total uptake of each root system with and without rhizodeposits for each rainfall distribution of the higher-rainfall regime.

| Total rainfall | 0.28 cm |  |  |  |  |  |
| --- | --- | --- | --- | --- | --- | --- |
| Rainfall distribution | 1 event |  | 2 events |  | 3 events |  |
| Rhizodeposits | No | Yes | No | Yes | No | Yes |
| Total uptake: 6-day old plant (cm) | $15.63 \times 10^{-3}$ | $13.65 \times 10^{-3}$ | $7.68 \times 10^{-3}$ | $9.15 \times 10^{-3}$ | $7.51 \times 10^{-3}$ | $9.18 \times 10^{-3}$ |
| Total uptake: 15-day old plant (cm) | $24.55 \times 10^{-3}$ | $23.21 \times 10^{-3}$ | $11.53 \times 10^{-3}$ | $14.99 \times 10^{-3}$ | $11.26 \times 10^{-3}$ | $15.02 \times 10^{-3}$ |
| Total uptake: 30-day old plant (cm) | $21.25 \times 10^{-3}$ | $27.95 \times 10^{-3}$ | $9.40 \times 10^{-3}$ | $15.55 \times 10^{-3}$ | $9.27 \times 10^{-3}$ | $15.62 \times 10^{-3}$ |

**Figure 7:**
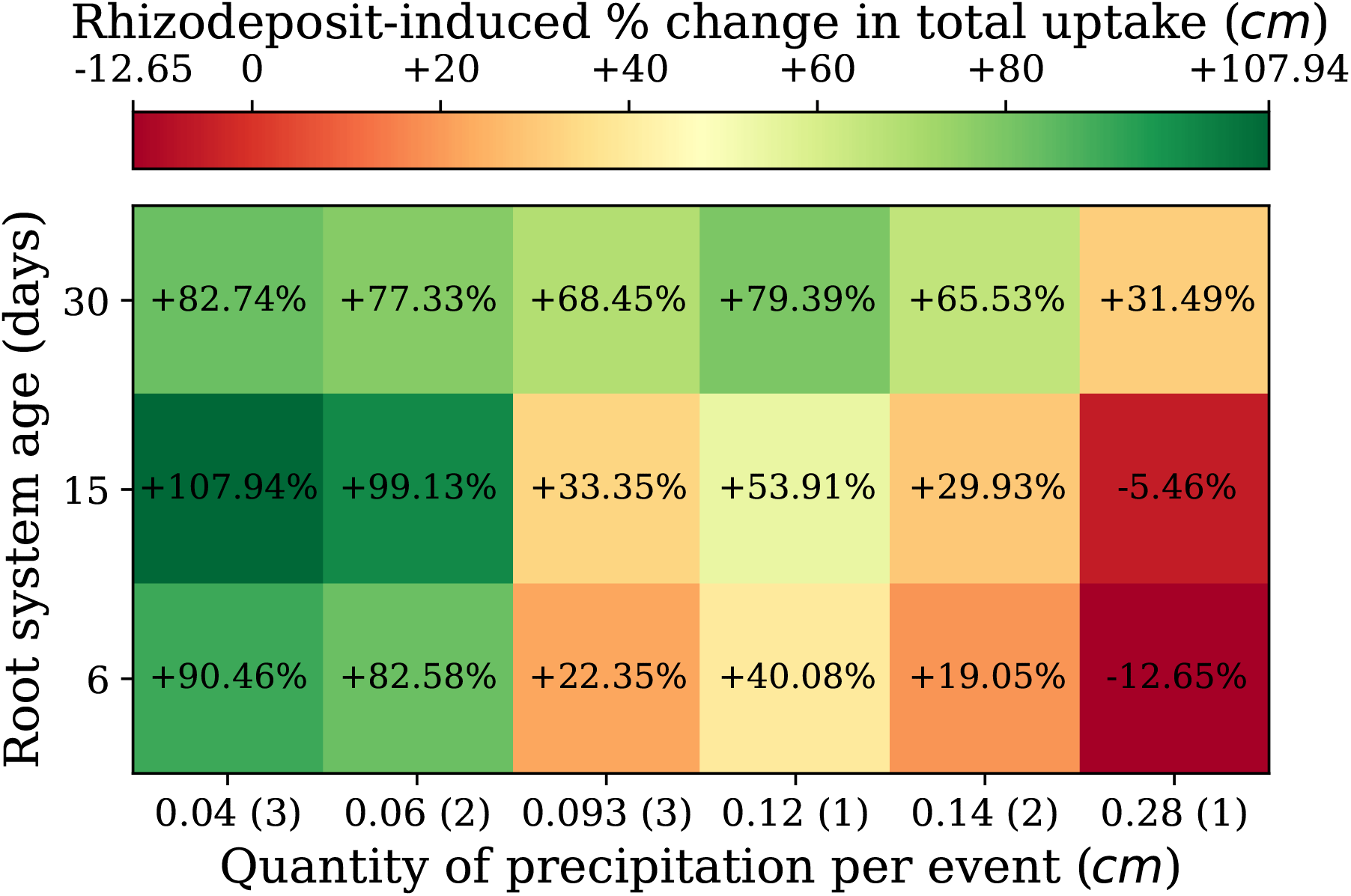
The strength of the positive or negative effect of rhizodeposits on total water uptake of a root system in relation to root system age and the quantity of precipitation delivered in each rainfall event of the precipitation regime. On the x-axis, numbers outside of brackets indicate the quantity of rainfall in one event of the regime. Numbers in brackets show the total number of events in that regime.

Regarding the increased uptake rates of the 15-day old root system compared to the 30-day old root system, which were observed during certain intervals of specific precipitation scenarios, the net effect on total uptake can also be seen in Tables 2 and 3. In the end, there were only 4 scenarios where the 15-day old root system had greater total uptake than the 30-day old root system. Namely, when total precipitation was 0.12 cm and all delivered in a single event with no rhizodeposits in the soil, when total precipitation was 0.12 cm and delivered over 2 or 3 events with rhizodeposits in the soil, and for all scenarios with 0.28 cm total precipitation and no rhizodeposits in the soil.

## 4 Discussion

### 4.1 Future incorporation of rhizodeposits and root water uptake in models of soil hydraulics

Within the field of mathematical modelling, there are different approaches to address rhizodeposit influence on soil hydraulic properties. Like in (Vogel et al., 1996; Karagunduz et al., 2001), the model (2.1)-(2.3) developed in this paper assumes that rhizodeposit-induced changes to contact angle and surface tension can be incorporated into the water retention (2.5) and hydraulic conductivity (2.6) functions of Mualem (1976) and Van Genuchten (1980) by using inverse air entry pressure and saturated hydraulic conductivity terms (2.13), (2.14) whose values are appropriately scaled according to the abundance of rhizodeposits. Similarly, Landl et al. (2021) described mucilage effects on hydraulics by adding a shift term to the Mualem (1976) and Van Genuchten (1980) water retention curve, which depended on mucilage and rhizosphere bulk density, and by scaling the saturated hydraulic conductivity with a term that depended on the viscosities of water and mucilage (Kroener et al., 2014). This was not the case, however, in (Cooper et al., 2017, 2018), where the influence of root exudates was incorporated into Richards equation through a rigorous multi-scale derivation, and the effect of increasing contact angle resulted in retention and conductivity curves that did not resemble the commonly used functions of Brooks and Corey (1964) or Mualem (1976) and Van Genuchten (1980).

As with the influence of rhizodeposits, a range of different approaches have also been taken to account for root water uptake when modelling soil water dynamics. At the largest scale there are the soil-plant-atmosphere continuum models (SPAC), which incorporate the physiological cycle of vegetation within simulations of water and energy dynamics at a landscape scale (Daly et al., 2004; Fatichi et al., 2016; Mencuccini et al., 2019). These are generally comprised of interlinked model compartments for processes such as radiation, fluxes of moisture and heat, plant respiration, and photosynthesis (Ivanov et al., 2008; Williams et al., 1996). For feasible simulation of these connected large-scale dynamics SPAC models tend to treat the individual processes in as simple a manner as possible, and the common model for root mass distribution is a function that simply decreases exponentially with depth. The contribution to transpiration of each soil layer is then based on its water content and the fraction of total root mass it contains (Fatichi et al., 2012; Feddes et al., 2001).

At the other end of the modelling spectrum are the approaches defined as mechanistic or microscopic, which more comprehensively account for a root system’s structure and hydraulic architecture. For example, below-ground water transport can be modelled in a way that gives explicit consideration to water flow through subdomains of soil and root tissue, with uptake driven by the difference in potential across the root-soil boundary (Roose and Fowler, 2004). In a similar spirit, water transport through a given root system has also been modelled by considering its hydraulic architecture as a network of connected nodes (Doussan et al., 1998; Javaux et al., 2008; Somma et al., 1998). In this framework, water moves down gradients in potential, with the axial and radial root segment conductivities determining the rates of water transport between roots and across the root-soil interface respectively (Couvreur et al., 2012). This results in a system of nonlinear equations that can be expressed in matrix form and solved to obtain the root water uptake rate at each root node. It should be noted, though, that this matrix system must be solved each time that the hydraulic heads of the soil, root nodes, or root collar change i.e. at every iteration of any coupled soil water transport model (Vanderborght et al., 2021). It is therefore the case with these mechanistic uptake models that as root system complexity and scale is increased the computational demand of obtaining simulations becomes prohibitively high.

The need for a compromise between the relatively basic approach to modelling uptake within SPAC systems and the mathematical complexity of mechanistic models has seen the introduction of empirical macroscopic root water uptake functions (Simunek and Hopmans, 2009), and one of these was employed in model (2.1). As shown in (2.15) macroscopic uptake functions *S* are generally expressed as a product of 4 terms: a piecewise-linear stress function *α*_*f*_ that decreases from 1 to 0 with decreasing pressure head, a normalised root length density function *NRLD* indicating the fraction of total root length within a given soil region, the potential maximum transpiration of the plant *T*_p_, and a compensation term (max (*ω, ω*_*c*_))^−1^, where *ω* is the *NRLD*-weighted average of the stress *α*_*f*_, and *ω*_*c*_ the critical water stress value, below which a lack of water in one region cannot be compensated through increased uptake by roots at other depths.

The benefit of these macroscopic models is that they provide an uptake sink term in Richards equation that is sensitive to local soil water content and incorporates compensatory uptake, while avoiding the computational demands of explicitly modelling flow into and through an entire root system.

There are, nevertheless, known disadvantages to macroscopic uptake functions. The first is that their parameters represent high level concepts, e.g. critical water stress *ω*_*c*_, and, hence, are more difficult to measure experimentally than those of mechanistic models (Cai et al., 2018). Secondly, assuming two regions of soil have identical levels of root density, and if, according to *α*_*f*_, neither of the two regions have water contents sufficiently low to constitute a stress to the roots, then the resulting uptake rate from these regions will be equal, despite the water contents potentially being considerably different (Vanderborght et al., 2024). Moreover, because they do not account for water potential within the root tissue, traditional macroscopic functions cannot capture hydraulic lift (Javaux et al., 2013). The results of this work, however, have shown another potential flaw in macroscopic uptake models such as (2.15) and that of Simunek and Hopmans (2009). The issue arises from the normalised root length density function in (2.15), which is used to determine a soil region’s maximum possible contribution to total uptake and computed by dividing the root length density function by the system’s total root length. This formulation means that as a root system grows larger, and the roots within a given region make up a smaller proportion of total root length, the value of *NRLD* for that region decreases (see Figures 3 and 4). As a result, even if the water content of a given soil layer and the conductance of the root system remain constant, the simulated uptake from the roots in that layer may well decrease as the root system grows. The consequence of this is then that a younger root system may exhibit higher rates and cumulative totals of simulated uptake than a mature root system, despite the younger plant having a much lower total root length within the wet regions of soil (Figures 5 and 6 and Tables 2 and 3).

In response to the problems regarding parametrisation and the modelling of hydraulic lift and water stress, Vanderborght et al. (2024) have recently proposed an alternative formulation of a macroscopic uptake function. In their approach the root system is again represented in a network format (Couvreur et al., 2012; Doussan et al., 1998; Javaux et al., 2008; Somma et al., 1998) but, instead of solving the resulting linear system, it is used to determine the contribution of each root to total root system conductance. These contributions, along with the transpiration demand, root collar water potential, and local soil water potential, are then used to determine the uptake from each soil layer. Furthermore, by applying homogenisation theory to a system of coupled equations for root and soil water transport, Mair and Ptashnyk (2024) recently derived a macroscopic equation for soil water transport where the term for root water uptake incorporates microscopic information like local hydraulic conductance and xylem water potential. In future, this could also be a viable method of formulating a macroscopic uptake function that is free from the issues raised previously.

### 4.2 Rhizodeposits as a mechanism for facilitated water infiltration

Many studies have found that the presence of roots in soil facilitates water infiltration following precipitation (Cerda, 1999; Marshall et al., 2014; Wu et al., 2016). Indeed, for a range of soil types, in comparison to being left fallow, their hydraulic conductivities have been found to be higher when vegetated by crops such as maize, willow and grass. (Feki et al., 2018; Leung et al., 2018). One of the proposed mechanisms behind this is the preferential flow of water through macropores that are created as growing roots, and the fauna attracted by their by-products, fragment and rearrange the soil (Angers and Caron, 1998; Luo et al., 2019). Additionally, rhizodeposits from roots have also been shown to have a surfactant effect when present in the soil water solution (Read and Gregory, 1997; Read et al., 2003; Naveed et al., 2019), and surfactants are often applied in agriculture to aid water infiltration into hydrophobic soils (Ogunmokun et al., 2020). It is, therefore, plausible that the facilitated water infiltration observed in vegetated soil can be partly attributed to the presence of certain rhizodeposits. Such an effect is captured within the formulation of our water transport model (2.1), where rhizodeposit-induced reductions to surface tension increase the soil hydraulic conductivity and alter the water retention curve so that infiltration of soil water is facilitated.

In our simulations where rhizodeposits were present, water infiltrated more quickly after the onset of precipitation and the rate of water loss by evaporation was reduced. This concurs with similar experimental results regarding evaporation losses under the application of bio-surfactants (Gutierrez et al., 2022). However, our results also highlighted that in terms of water availability to the root system under enhanced infiltration, a clear trade off exists between reduced evaporation losses and increased deep percolation. Furthermore, the net effect of this trade off will vary with soil type, precipitation pattern and root system morphology. In very conductive sandy soils, where water retention is comparatively poor (Bhardwaj et al., 2007), low irrigation rates tend to be recommended for optimal root water uptake efficiency (Alhammadi and Al-Shrouf, 2013) since increases to infiltration rate only reduce the availability of water to roots. In contrast, for less conductive soils water losses from evaporation are likely to be larger than from deep percolation and, therefore, increases to infiltration rate improve root water uptake and crop yield (Amooh and Bonsu, 2015; Oostindie et al., 2010). In terms of precipitation pattern, Chaichi et al. (2015) found that if rainfall or irrigation events were smaller and total water was delivered more gradually, then the effect of enhanced infiltration on root water uptake and yield was positive. But, if total rainfall is concentrated into fewer but more intense events, then the enhancement of downward flow can become less beneficial or even detrimental to the availability of water to the root system (Blackwell, 2000). These findings were borne out by our results, with Figure 7 showing for root systems of all ages that rhizodeposit-induced increases to infiltration were generally less beneficial to total root water uptake as the quantity of rainfall per event increased. Moreover, when all 0.28 cm of precipitation was delivered in one rainfall event, the total uptake of the 6 and 15-day old root systems was actually worsened by the presence of rhizodeposits (Figure 7). The intuitive explanation for this is that infiltration becomes too strong. That is, despite the water’s arrival to the root zone being accelerated and initially improving uptake rates (Figure 6), ultimately its faster infiltration means that the window of time it is available for uptake by these shallower root systems is shorter than when rhizodeposits are not present (Figure 4). In contrast, the deeper roots of the 30-day old root system mean that the rhizodeposit-induced increases to infiltration never negatively affected uptake, regardless of precipitation pattern (Figure 7). Such a result concurs with the experimental findings of Shao et al. (2009) where, due to their larger root systems, the water uptake efficiency of mature wheat root systems appeared robust under alterations to irrigation pattern.

In summary, the findings of our work suggest that the consequences on root water uptake of infiltration-boosting rhizodeposits depends upon the environmental conditions considered and is not necessarily universally positive. The important qualification to accompany our conclusions is that not all rhizodeposits influence soil hydraulics in the same way. For example, there is evidence of maize, barley and chia increasing the viscosity of soil water solutions (Naveed et al., 2019), which would reduce hydraulic conductivity in the rhizosphere and hinder the movement of water around the root. In addition to this, the characteristics of the rhizodeposits of a given plant species are also almost certainly not constant over time. In fact, there is growing evidence that the production rate and chemical profile of rhizodeposits are adaptive to biotic factors like growth stage, and external factors such as soil type and water or nutrient availability (Rolfe et al., 2019; Wen et al., 2022; Williams and de Vries, 2020). This, therefore, gives weight to claims that rhizodeposits are deployed by a plant to engineer the soil in accordance to their needs, as opposed to being merely a waste product whose potential effects on water and nutrient availability are merely coincidental.

### 4.3 Harnessing rhizodeposits for the development of crop ideotypes with resilience to enviroment-specific stresses

The Green Revolution of the 1960s, with the increased use of chemical fertilisers, pesticides and precise irrigation, shifted the attention of crop breeders towards the question of maximising yield under conditions where stresses from water and nutrient shortages or competition were minimal (Preece and Peñuelas, 2020). This approach led to a focus on optimal above ground traits, where one direction was to breed for crops that maximised the ratio of harvestable above ground biomass to total above ground biomass (the harvest index) (Richards et al., 1993). Since progress in this aspect rapidly became marginal, improvements to total above ground biomass were then sought, and achieved, through breeding for rapid leaf area development to reduce evaporation losses from the soil and enhance leaf-level water use efficiency. Within the context of roots, however, the aim of many was to maintain root development at the necessary minimum so as to maximise resource allocation to harvestable biomass under ideal growing conditions (Condon et al., 2004). Nevertheless, with the increasing incidence of global drought, breeding strategies for resilient crops have latterly started to consider more developed root systems with features that improve access to soil water.

Genes have been identified that control root gravitropic response and radial growth (Hafeez et al., 2024) and there is a belief that increasing rooting depth should be the target of breeders. This is firstly because it allows rain-fed crops to access deep stores of water during periods of drought (Lynch, 2013; Uga et al., 2013) and, secondly, because in wheat systems root arrival at deep water stores tends to coincide with grain-filling season thus boosting pre-harvest biomass (Palta et al., 2011; Wasson et al., 2012; Ober et al., 2021). Evidence has also been shown of a genome effect on rhizodeposit and microbiome features (Iannucci et al., 2021), and it has been proposed that crops could be bred to promote the development of microbiomes that are specifically beneficial under certain stresses (Ober et al., 2021). For instance, as suggested by the findings of this work, if crop root systems are deep and soil water losses are dominated by evaporation, then the production of rhizodeposits with surfactant like properties could improve root water uptake.

Despite this potential, however, few crop breeding programs currently incorporate rhizodeposit characteristics in there selection criteria (Preece and Peñuelas, 2020). One reason for this is that it is far from straightforward to verify that a given crop genotype produces rhizodeposits that affect soil characteristics in the way that they have been purported to. This is because non-destructively extracting rhizodeposits from roots grown in the field is massively challenging, and when taking the more simple approach of extracting from roots grown in hydroponic systems, it cannot be guaranteed that the same rhizodeposit properties would be observed for roots grown in soil (Oburger and Jones, 2018). Nevertheless, increasingly sophisticated methods are now being employed to overcome these hurdles (Casas and Matamoros, 2021) such as growing roots in soil then transferring to hydroponic systems for rhizodeposit extraction (Gargallo-Garriga et al., 2018; Herz et al., 2018), sampling rhizodeposits directly from soil by leachate collection (de la Paz et al., 2019), and the use of matrix-assisted laser ionization for more rapid mass-spectrometry analysis (Sasse et al., 2020). With these technological advances, knowledge of rhizodeposit properties continues to improve. Additionally, the emergence of synthetic soil microcosm systems is facilitating the observation of the effects that these rhizodeposits have on soil hydraulic processes at the microscopic level (Hayat et al., 2021; Liu et al., 2025; Zarebanadkouki et al., 2019). Continued progress in this area will generate data that allows the improved formulation and calibration of models (like the one presented in this work) so that they better represent the impact of rhizodeposits on soil hydraulics in both a qualitative and quantitative sense. In time, such models could help to predict the consequences of given rhizodeposits on macroscopic metrics of crop performance like water use efficiency and yield. None of these steps are by any means trivial, but their completion should make the consideration of rhizodeposit properties within crop breeding programs a more feasible prospect.

## Acknowledgements

This work was supported by the European Union’s Horizon Europe under the grant agreement No.101060124 (Project Root2Res). We also acknowledge the funding from the Spanish Ministry of Science, Innovation and Universities under the projects MICROCROWD and BIOFLOW (PID2020-112950RR-I00, PID2023-149435OR-I00).

## Data availability statement

The data generated by all simulations, along with the codes to parametrise and numerically solve the model equations, are available from Zenodo via https://doi.org/10.5281/zenodo.15101432 under a Creative Commons Attribution 4.0 International license.

